# Generative artificial intelligence performs rudimentary structural biology modeling

**DOI:** 10.1101/2024.01.10.575113

**Authors:** Alexander M. Ille, Christopher Markosian, Stephen K. Burley, Michael B. Mathews, Renata Pasqualini, Wadih Arap

## Abstract

Natural language-based generative artificial intelligence (AI) has become increasingly prevalent in scientific research. Intriguingly, capabilities of generative pre-trained transformer (GPT) language models beyond the scope of natural language tasks have recently been identified. Here we explored how GPT-4 might be able to perform rudimentary structural biology modeling. We prompted GPT-4 to model 3D structures for the 20 standard amino acids and an α-helical polypeptide chain, with the latter incorporating Wolfram mathematical computation. We also used GPT-4 to perform structural interaction analysis between nirmatrelvir and its target, the SARS-CoV-2 main protease. Geometric parameters of the generated structures typically approximated close to experimental references. However, modeling was sporadically error-prone and molecular complexity was not well tolerated. Interaction analysis further revealed the ability of GPT-4 to identify specific amino acid residues involved in ligand binding along with corresponding bond distances. Despite current limitations, we show the capacity of natural language generative AI to perform basic structural biology modeling and interaction analysis with atomic-scale accuracy.

## Introduction

Artificial intelligence (AI)-based capabilities and applications in scientific research have made remarkable progress over the past few years [1, 2]. Advances in the field of protein structure prediction have been particularly impactful: AI-based dedicated structural biology tools such as AlphaFold2 and RoseTTAFold are capable of processing amino acid residue sequences only as input and modeling their corresponding protein structures with accuracy comparable to lower-resolution experimentally determined structures [3-5]. AlphaFold2 and RoseTTAFold are trained on datasets of protein sequence and structure datasets [6], and rely on neural network architectures specialized for modeling protein structures. Another category of AI-based tools for protein structure prediction are protein language models, which differ from AlphaFold2 and RoseTTAFold in that they are not trained on structures but rather on protein sequences [7-9]. Collectively, such protein structure prediction tools have been extensively used by researchers across various disciplines in the biological sciences and are expected to continue to add value alongside experimental structure determination [10-18].

On the other hand, generative AI language models, particularly the various generative pre-trained transformer (GPT) models powering ChatGPT [19-21], have garnered substantial interest with applications including for performance of autonomous and predictive chemical research [22, 23], drug discovery [24-26], bioinformatic analysis [27-30], synthetic biology [31], and the potential to aid in clinical diagnosis [32, 33], among others [34-36]. Unlike AlphaFold2, RoseTTAFold, and protein language models, the GPT family of language models [19, 20] are trained on natural language datasets and its neural network architecture is tailored towards understanding and generating natural language text rather than structural modeling. Our group has recently reported how GPT-4 interprets the central dogma of molecular biology and the genetic code [37]. While analogies can be made, the genetic code is not a natural language *per se*, yet GPT-4 seems to have an inherent capability of processing it. In a similar line of investigation, here we explored whether GPT-4 could perform rudimentary structural biology modeling and evaluate its capabilities and limitations in this domain. Surprisingly, we found that GPT-4 is indeed capable of modeling the 20 standard amino acids and, with incorporation of the Wolfram plugin, a typical α-helical secondary structure element at the atomic level—albeit with some errors. Moreover, we used GPT-4 to perform structural analysis of interaction between the drug nirmatrelvir and its molecular target, the main protease of SARS-CoV-2 (the coronaviral etiology of COVID-19). To our knowledge, this is the first report to explore the specific capabilities of GPT-4 in rudimentary structural biology modeling and protein-ligand interaction analysis.

## Results

### Modeling of individual amino acid structures

Amino acid residues are the components of proteins, and their atomic composition and geometric parameters have been well characterized [38-40], making them suitable candidates for rudimentary structure modeling. We therefore prompted GPT-4 to model the 20 standard amino acids with minimal contextual information as input, including instructions for output in legacy Protein Data Bank (PDB) file format **(Table 1a, 2a, S1, and Fig. 1a)**. GPT-3.5 was included as a performance benchmark. Multiple iterations (n=5 for each amino acid) were run by using the same input prompt to monitor consistency (see Methods). For each individual amino acid, GPT-4 generated 3D structures with coordinate values for both backbone and sidechain atoms **(Fig. 1b)**. Generated structures contained all atoms specific to the amino acid prompted, except for a single iteration of cysteine which lacked the backbone O atom and a single iteration of methionine which lacked sidechain Cγ atom. Most amino acid structures (excluding achiral glycine) were modeled in L rather than D stereochemical configuration, while some were also modeled in planar configuration **(Fig. 1c)**. While the modeling favored the L-configuration, a more accurate distribution would be near exclusive L-configuration, given that D-amino acid residues are only rarely found in naturally occurring proteins [41].

**Table 1.** Prompts used for structural modeling. **(A)** Prompt used for modeling the structures of each of the 20 amino acids with GPT-4 and GPT-3.5. The same prompt was used for each amino acid by replacing “[amino acid]” with the full individual amino acid name. **(B)** Prompt used for modeling the α-helical polypeptide structure with GPT-4 running with the Wolfram plugin for enhanced mathematical computation. **(C)** Prompts used for structural drug interaction analysis of nirmatrelvir bound to the SARS-CoV-2 main protease (PDB ID: 7VH8).

|  |  |
| --- | --- |
| **(A) Amino acid structure modeling with GPT-4 and GPT-3.5** |  |
| **Prompt:** What are the typical distances and angles between the atoms of one [amino acid] residue in a protein? Based on these values, generate a structure in PDB file format for one [amino acid] residue. Ensure coordinate values have three decimal places and omit hydrogen atoms. |  |
| **(B) α-helix structure modeling with GPT-4 running the Wolfram plugin** |  |
| **Initial Prompt:** What are the typical geometric attributes (including distances and angles) between the backbone atoms of a typical alpha-helical polypeptide chain? Based on this information, generate a structure in PDB file format for an alpha-helical polypeptide chain 10 residues in length including only alpha carbon atoms. Ensure coordinate values have 3 decimal places and use the Wolfram plug-in for coordinate calculations but not for PDB file formatting. |  |
| **Refinement Prompts** | The generated structure does not resemble an alpha-helix. Try again. |
|  | The diameter of the helix is too large. |
|  | The diameter of the helix is too small. |
|  | The pitch (residues per turn) of the helix is too large. |
|  | The pitch (residues per turn) of the helix is too small. |
| **(C) Structural drug interaction analysis with GPT-4** |  |
| **Ligand detection prompt:** | Based on structural information, what is the ligand that is present in the attached PDB file? |
| **Interaction detection prompt:** | Analyze the structure to detect up to 5 amino acid residues in the protein chain which have important bond interactions with the "4WI" ligand without importing |
|  | any external libraries. List the residues and the bond distances. Based on this information, predict potential mutations in the protein chain which would interfere with binding of the "4WI" ligand. |

**Fig. 1.**
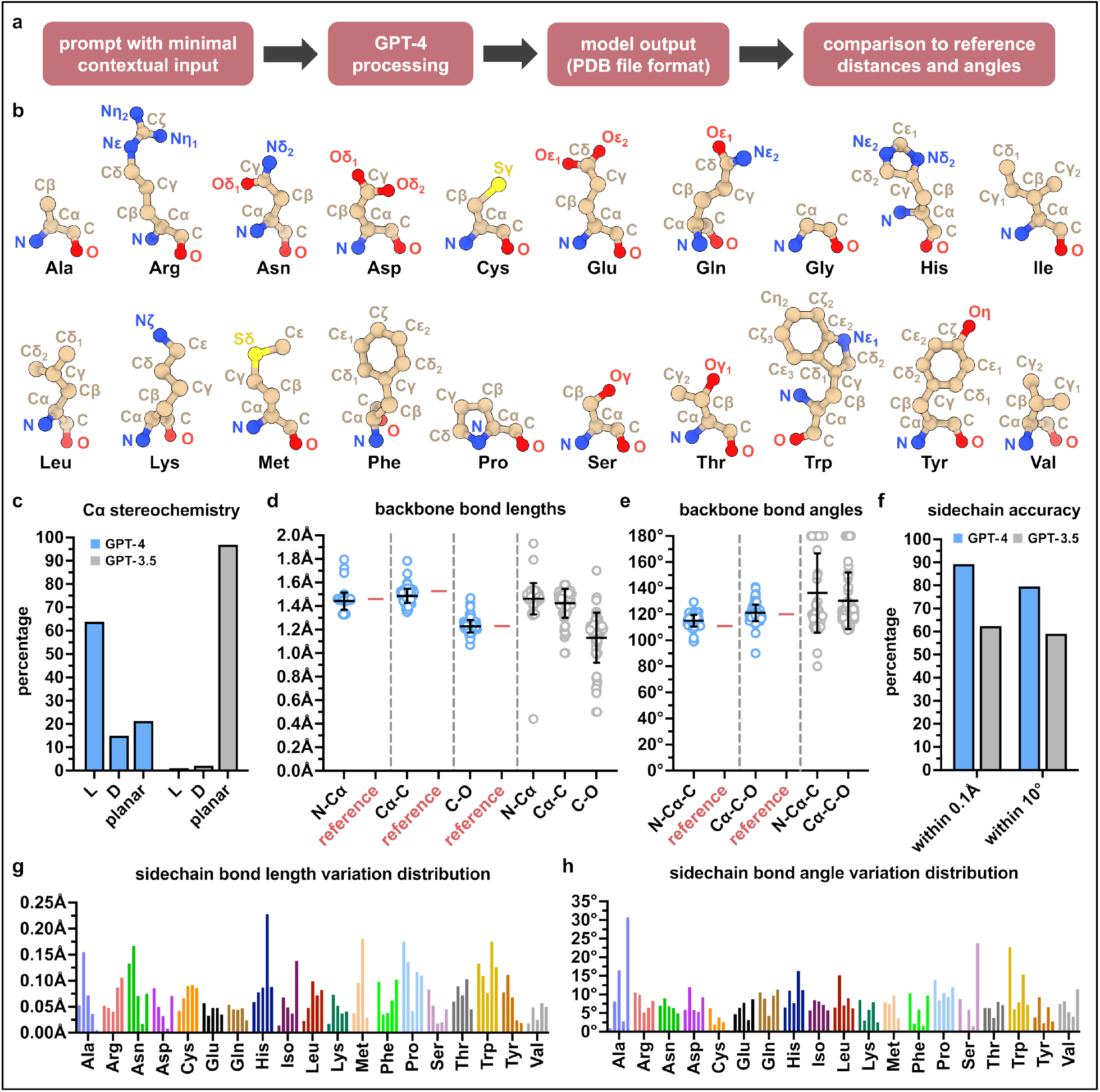
Modeling the 3D structures of the 20 standard amino acids with GPT-4. **(a)** Procedure for structure modeling and analysis. **(b)** Exemplary 3D structures of each of the 20 amino acids modeled by GPT-4. **(c)** Cα stereochemistry of modeled amino acids including L and D configurations as well as nonconforming planar, n=5 per amino acid excluding achiral glycine and one GPT-4 iteration of cysteine (see methods). **(d-e)** Backbone bond lengths and angles of amino acids modeled by GPT-4 (blue) relative to experimentally determined reference values (red), n=5 per amino acid, excluding one iteration of cysteine (see methods). Corresponding values of amino acids modeled by GPT-3.5 shown adjacent (grey), n=5 per amino acid. Data shown as means ± SD. **(f)** Sidechain accuracy of modeled amino acid structures in terms of bond lengths (within 0.1 Å) and bond angles (within 10°) relative to experimentally determined reference values, n=5 per amino acid. See Methods for experimentally determined references. **(g-h)** Distributions of sidechain bond length and angle variation relative to experimentally determined reference values for each amino acid generated by GPT-4, excluding glycine. Bars represent the mean bond length or angle variation for each of the five iterations per amino acid. One of the methionine iterations was excluded (see methods).

Backbone bond lengths and angles of the modeled structures varied in accuracy, yet clustered in approximation to experimentally determined reference values [38, 39] **(Fig. 1, d and e)**. Moreover, all reference values fell within the standard deviations of backbone bond lengths and angles of the modeled structures. Finally, sidechain bond lengths and bond angles also varied in accuracy, yet >89% of calculated bond lengths were within 0.1 Å and >79% of calculated bond angles were within 10° of experimentally determined reference values [40, 42], again indicating remarkable precision **(Fig. 1f-h, Fig. S1-S4)**. Sidechain bond lengths and bond angles outside of these ranges generally occurred at random, but were notably more prevalent in the aromatic rings of histidine and tryptophan, along with the pyrrolidine component of proline. Although not entirely error-free, the ring structures of phenylalanine and tyrosine were more accurate, which may be due to the reduced complexity of their all-carbon ring composition. Across all parameters assessed, GPT-4 substantially outperformed GPT-3.5. Collectively, these findings demonstrate that GPT-4 is capable of structurally modeling single amino acid residues in a manner that resembles their experimentally-determined structures, though not without sporadic errors including incorrect stereochemistry and geometric distortion, which would require—at least presently—human operator curation or supervision to ensure fidelity.

### Modeling of an α-helix structure

The α-helix is the most commonly occurring and extensively studied secondary structure element found in proteins [43-46]. Thus, we next prompted GPT-4 and GPT-3.5 to model an α-helical polypeptide chain, but were unable to obtain accurate structures with either version, despite multiple attempts with various prompts. We then incorporated the Wolfram plugin, a mathematical computation extension developed by Wolfram-Alpha for use with GPT-4 [47]. GPT-4 used together with the Wolfram plugin was able to model a 10-residue α-helical structure and output the result in legacy PDB file format with minimal contextual information as input **(Table 1b, 2b, S2, and Fig. 2a)**. Multiple iterations were run by using the same input prompt to monitor consistency, and up to two prompt-based refinements after the first attempt were permitted per iteration for improved accuracy (see Methods). To reduce complexity, only Cα atoms were modeled.

**Table 2.**
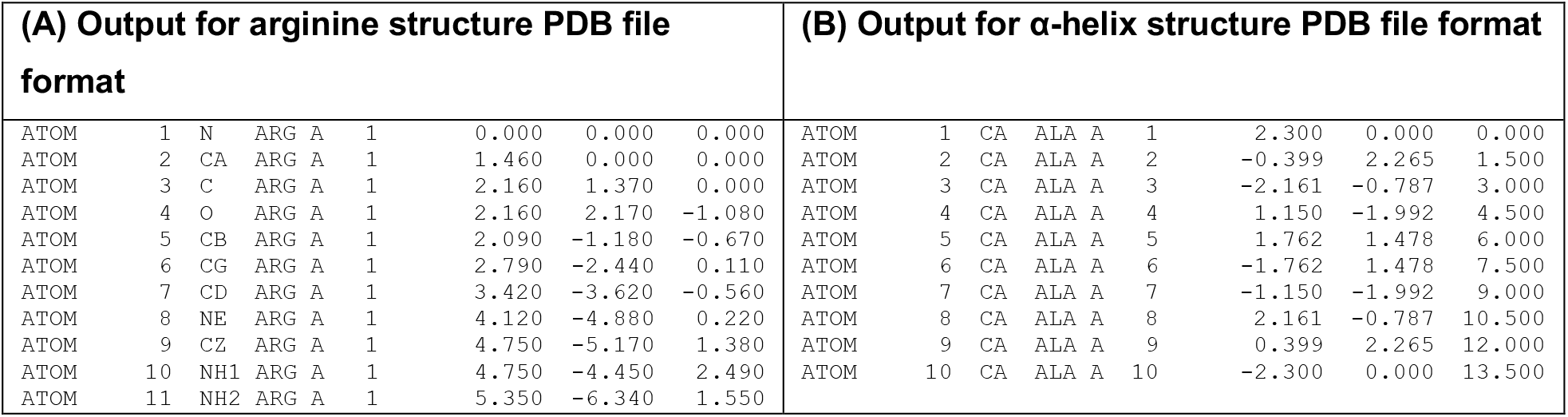
Exemplary coordinate output from GPT-4. Within responses to modeling prompts **(Table 2, S1, and S2)** GPT-4 provided coordinates for generated structures in PDB file format, as shown below with output examples for **(A)** the arginine amino acid structure and **(B)** the α-helix structure.

**Fig. 2.**
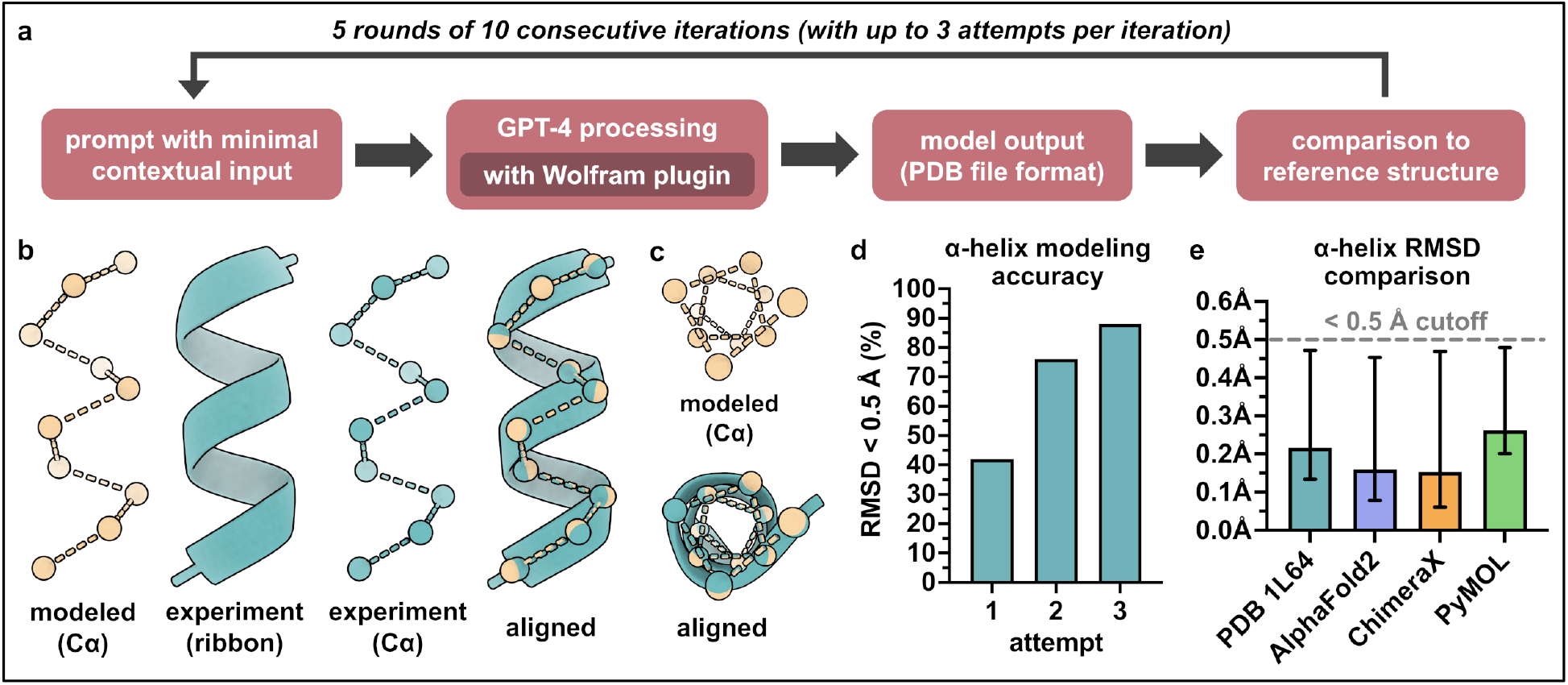
Modeling the 3D structure of an α-helical polypeptide structure with GPT-4. **(a)** Procedure for structure modeling and analysis. **(b)** Exemplary 3D structure of a modeled α-helix (beige), an experimentally determined α-helix reference structure (PDB ID 1L64) (teal), and their alignment (RMSD = 0.147 Å). **(c)** Top-down view of modeled and experimental α-helices from panel B. **(d)** Accuracy of α-helix modeling as measured by number of attempts (including up to two refinements following the first attempt) required to generate a structure with RMSD < 0.5 Å relative to the experimentally determined reference structure, n = 5 rounds of 10 consecutive iterations (total n = 50 models). **(e)** Comparison of RMSDs between GPT-4 α-helix structures and the experimentally determined α-helix structure, the AlphaFold2 α-helix structure, the ChimeraX α-helix structure, and the PyMOL α-helix structure. Only structures with RMSD < 0.5 Å (dashed grey line) relative to each reference structure are included (88% included in reference to PDB ID 1L64; 90% to AlphaFold; 90% to ChimeraX; 88% to PyMOL). Data shown as means ± range.

GPT-4 arbitrarily assigned all residues as alanine, which was likely done for the sake of simplicity, but nevertheless aligns well with the fact that alanine has the greatest α-helix propensity of all 20 standard amino acids [46]. Remarkably, accuracy of the modeled α-helix was comparable to an experimentally determined α-helical structure consisting of 10 consecutive alanine residues (PDB ID: 1L64) [48] **(Fig. 2, b and c)**. Forty-four percent of modeled structures had a root-mean-square deviation (RMSD) of < 0.5 Å relative to the reference experimental structure on the first attempt, and 88% had a RMSD of < 0.5 Å after two prompt-based refinements **(Fig. 2d)**. The structures generated by GPT-4 were also compared to poly-alanine α-helix structures modeled by AlphaFold2, ChimeraX, and PyMOL, and the lowest RMSDs (*i*.*e*., greatest structural similarity) were found between the AlphaFold2 and ChimeraX structures **(Fig. 2b)**. Taken together, these results demonstrate the capability of GPT-4, with a seamless incorporation of the Wolfram plugin, to predict the atomic level structure of an α-helix.

### Structural interaction analysis

Structural interaction between drugs and proteins is a key aspect of molecular biology with basic, translational and clinical implications. For instance, binding of the Paxlovid (ritonavir-boosted nirmatrelvir) protease inhibitor compound, nirmatrelvir, to the SARS-CoV-2 main protease is of particular clinical relevance [49, 50], especially given the concern that the mutation-prone SARS-CoV-2 leads to treatment resistance [51]. Thus, we used GPT-4 to perform qualitative structural analysis of drug binding within the nirmatrelvir-SARS-CoV-2 drug-protein paradigm. We first provided the PDB file input of a crystal structure of nirmatrelvir bound to the SARS-CoV-2 main protease (PDB ID: 7VH8) [49] and prompted GPT-4 to detect the nirmatrevir ligand, followed by a subsequent prompt for interaction detection and interaction-interfering mutation prediction **(Table 1c, S3, and Fig. 3a)**.

**Fig. 3.**
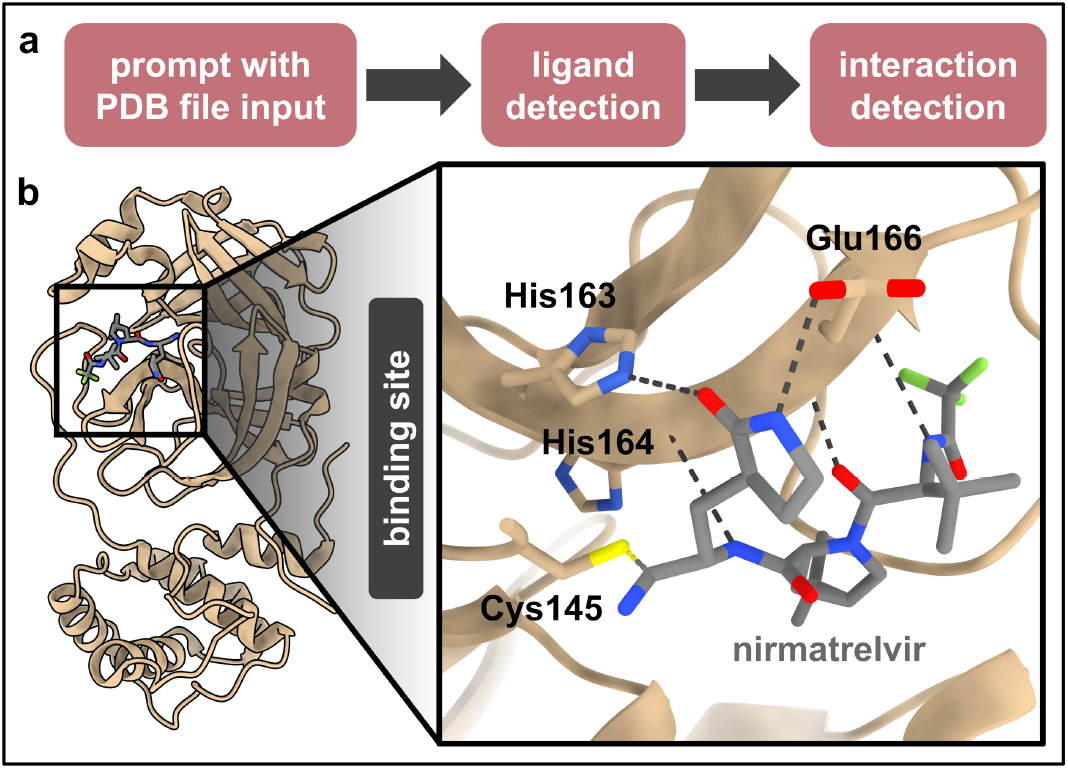
Structural analysis of interaction between nirmatrelvir and the SARS-CoV-2 main protease. **(a)** Procedure for performing ligand interaction analysis. **(b)** Crystal structure of nirmatrelvir bound to the SARS-CoV-2 main protease (PDB ID: 7VH8) with bond-forming residues detected by GPT-4, and their bonds depicted with ChimeraX (inset). Distances between interacting atom pairs were 1.81Å (Cys145 Sγ– C3), 2.68 Å (His163 Nε2–O1), 2.77Å (Glu166 O–N4), 3.02Å (His164 O–N1), as determined by GPT-4 and 1.814Å (Cys145 Sγ–C3), 2.676 Å (His163 Nε2–O1), 2.767Å (Glu166 O–N4), 2.851Å (Glu166 N–O3), 3.019Å (Glu166 Oε1–N2), 3.017Å (His164 O–N1), as determined with ChimeraX. Note that distance values corresponding to the Glu166 N–O3 and Glu166 Oε1–N2 atom pair interactions were not provided by GPT-4.

GPT-4 correctly identified the nirmatrelvir ligand, which in the input PDB file is designated as “4WI” **(Table S3)**. For interaction detection, GPT-4 listed five amino acid residues within the substrate-binding pocket of the protein, four of which directly bind the nirmatrelvir ligand (Cys145 forms a covalent bond, His163 and His164 each form hydrogen bonds, and Glu166 forms three separate hydrogen bonds) [49] **(Fig. 3b)**. The fifth residue (Thr190) does not form a bond with the ligand, but is located within the binding pocket [49]. Moreover, the distances provided by GPT-4 for the four binding residues correspond precisely to the distances between the interacting atoms, information which is not inherent in the input PDB file. GPT-4 also described several mutations which may interfere with binding **(Table S3)**, and while most were plausible, others would likely be inconsequential. Notably, however, the suggested mutation of Glu166 to a residue lacking negative charge has been documented to be critically detrimental to nirmatrelvir binding [52-54] and confers clinical therapeutic resistance [55, 56]. Altogether, this exercise reveals the ability of GPT-4 to perform basic structural analysis of ligand interaction in a manner which, in conjunction with molecular analysis software such as ChimeraX, highlights its practical utility.

## Discussion

The proof-of-concept findings reported here show the current capabilities and limitations of GPT-4, a natural language-based generative AI, for rudimentary structural biology modeling and drug interaction analysis. This presents a unique aspect of novelty, given the inherent distinction between natural language models and other dedicated AI tools commonly used for structural biology, including AlphaFold, RosettaFold, and protein language models. While such tools are unequivocally far more advanced in terms of the scale of molecular complexity they are able to process, GPT-4 sets the stage for a broadly accessible and computationally distinct avenue for use in structural biology and protein-ligand interaction analysis.

The performance of GPT-4 for modeling of the 20 standard amino acids was favorable in terms of atom composition, bond lengths, and bond angles. However, stereochemical configuration propensity and modeling of ring structures require improvement. Performance for α-helix modeling, with a seamless incorporation of advanced mathematical computation from the Wolfram plugin, was also favorable. While the requirement of prompt-based refinements may be viewed as a limitation, they may also serve as a means and opportunity by which the user can optimize and modify a structure. Nonetheless, improvements will be required in the capacity to model more complex all-atom structures, not only Cα backbone atoms.

These structural modeling capabilities also raise the question of modeling methodology, especially since GPT-4 was not explicitly developed for this specialized purpose. It would be challenging to provide a precise answer for this, and several (proprietary or non-proprietary) computational methods may be involved. For instance, GPT-4 may be utilizing pre-existing atomic coordinate information present in its broad training dataset, which includes “publicly available data (such as internet data) and data licensed from third-party providers” [20]. However, this reasoning does not adequately explain the geometric variability observed in the predicted structures, and why structural complexity appears to be a limiting factor. The modeling may also be performed *ab initio*, given that the generated responses often articulate geometric parameters (*e*.*g*., specific bond lengths and angles, number of amino acid residues per α-helix turn, α-helix diameter, etc.) in addition to providing atomic coordinates **(Table S1 and S2)**. Alternatively, the modeling methodology may involve both the use of pre-existing coordinates plus *ab initio* computation.

Of note, the comparison of the α-helix model generated by GPT-4 with those generated by other computational tools was quite revealing. AlphaFold2, as mentioned above, predicts structures based on training data consisting of protein sequences and 3D structures, and was developed specifically for modeling protein structures. In addition to their dedicated molecular analysis capabilities, ChimeraX and PyMOL may be used to model basic, idealized secondary structure elements in a manner which narrowly considers precise predefined geometries, thus providing accurate α-helix structures. Despite not being explicitly developed to model atomic coordinates for α-helical segments of protein chains, GPT-4 was able to generate an α-helix with accuracy comparable to the structures modeled by the above tools. The requirement of adding the Wolfram plugin likely suggests that mathematical computation is heavily relied upon by GPT-4 for α-helix modeling. Yet, α-helix structural properties and self-instruction are generated by GPT-4 prior to engaging the Wolfram plugin **(Table S2)**, suggesting that some degree of intrinsic “reasoning” might perhaps be involved. So-called reasoning, in this regard, refers to the documented performance of benchmark reasoning exercises by GPTs [19, 20], and it should be noted that there is ongoing debate about what constitutes reasoning as it pertains to AI [57, 58]. While this work was under external peer review, a new version of GPT-4 (GPT-4o) was released by OpenAI, which we found to be capable of accurate α-helix modeling without the use of the Wolfram plugin. This new finding further demonstrates the capabilities and potential of structural modeling emerging from the rapidly evolving field of natural language-based generative AI.

The exercise exploring the capability of GPT-4 to perform structural analysis of ligand-protein binding showed promise, especially given the clinical relevance of the protein binding interaction between nirmatrelvir and the SARS-CoV-2 main protease. Ligand detection was expected to be a straightforward task, as PDB files include unique designations for various molecular entities. Interaction detection was surprisingly well-handled, considering the complexity of locating amino acid residues with spatial proximity to the ligand and providing precise distances between interacting atoms. Based on the generated response **(Table S3)**, it is likely that proximity was the primary criteria used by GPT-4 for interaction detection. While proximity is important, the analysis would benefit from additional criteria such as hydrophobicity, electrostatic potential, solvent effects, etc. These criteria would likely improve the prediction of interaction-interfering mutations.

Considering both strengths and weaknesses, the structural modeling capabilities of GPT represent an intriguing aspect of the unprecedented advancement of natural language-based generative AI, a transformative technology still in its infancy. While this modeling remains rudimentary and is currently of limited practical utility, it establishes an immediate and direct precedent for applying this technology in structural biology as generative AI natural language models undergo continued development and specialization. Concurrently, this broadly-accessible technology presents opportunity for structural analysis of drug-protein interaction. In the interim, further research on the capabilities and limitations of generative AI is merited, not only in structural biology but also for other potential applications in the biological sciences.

## Methods

### Prompt-based modeling with GPT-4

Modeling of individual amino acid structures was performed by challenging GPT-4 through the ChatGPT interface [20, 21] with a single prompt **(Table 1a)**, one amino acid residue at a time. For each individual amino acid, the same prompt was used for five consecutive iterations with each iteration initiated in a new dialog. GPT-4 was run in classic mode without “browser” and “analysis” features enabled, formerly known as “web browser” and “code interpreter” plug-ins, respectively [59]. Classic mode limits processing to GPT-4 with no additional capabilities. Amino acid modeling was also performed with GPT-3.5 in the same manner. However, GPT-3.5 would frequently generate PDB file output with missing or extra atoms. In such cases, responses were regenerated within each GPT-3.5 dialog until PDB file output contained the correct number of atoms required for analysis. Modeling of α-helix structures was performed by challenging GPT-4 running the Wolfram plugin [60] through the ChatGPT interface with an initial prompt followed by up to two refinement prompts in the same dialog, for a total of up to three attempts **(Table 1b)**. The same prompt was used for five rounds of ten consecutive iterations with each iteration initiated in a new dialog.

### Analysis of generated structures

Structures were analyzed by using the UCSF ChimeraX [42]. For amino acid structures, the “distance” (for bond lengths) and “angle” (for bond angles) commands were used with atom specification tailored to each amino acid type. Experimentally determined amino acid backbone bond lengths (N-Cα, 1.458Å; Cα-C, 1.525Å; C-O 1.231Å) [38], backbone bond angles (N-Cα-C, 111.0°; Cα-C-O, 120.1°) [39], and sidechain bond lengths and bond angles **(Table S4)** [40] served as references for evaluating predicted amino acid structures. Sidechain bond lengths and bond angles, obtained from a backbone-dependent rotamer library built into ChimeraX, represent those in which the backbone dihedral angles are ω=180° (typical of trans peptide bonds), φ=180°, and ψ=180° [40, 42]. For GPT-4, one iteration of cysteine lacked the backbone O atom and one iteration of methionine lacked the sidechain Cγ atom. Thus, these single iterations (n=1) from were excluded from analyses involving the missing atoms.

For α-helix structures, the matchmaker tool was used for alignment and RMSD determination across all atom pairs by using its default parameters (*i*.*e*., best-aligning pair). An experimentally determined α-helical structure consisting of 10 consecutive alanine residues, detected within an engineered form of bacteriophage T4 lysozyme (PDB ID 1L64) [48], was used as the reference for evaluating the modeled α-helix structures. The AlphaFold2 α-helix structure was modeled using ColabFold [61] through ChimeraX run with default parameters (*i*.*e*., minimization and without templates usage) using an elongated polyalanine sequence as input **(Table S5 and Fig. S5a)**. The ChimeraX and PyMOL (version 2.5.7) α-helix structures were modeled by using the build structure command and fab command, respectively, each by using a 10-residue alanine sequence as input and run with default α-helix parameters (*i*.*e*., backbone dihedral angles set to φ=-57° and ψ=-47°) **(Fig. S5, b and c)**. All data were analyzed using GraphPad Prism 10.1.0 (GraphPad Software). Statistical details are reported in the figure legends and statistical measurements include mean, mean ± SD, and mean ± range.

### Prompt-based interaction analysis with GPT-4

Structural analysis of binding interaction was performed by providing GPT-4 with an input PDB file and prompting as described (Table 1B) through the ChatGPT interface. The PDB file used as input was unmodified as obtained from the PDB entry for PDB ID: 7VH8 [49]. It should be noted that PDB ID: 7VH8 refers to nirmatrelvir as PF-07321332. ChimeraX was used to analyze amino acid residues detected by GPT-4 to interact with nirmatrelvir. The “contacts” and “distance” commands were used to verify interactions. For this exercise, GPT-4 was not limited to classic mode. Rather the “browser” and “analysis” features were enabled within the ChatGPT interface to enable file input, a feature available for GPT-4 but not GPT-3.5. Only the “analysis” feature was engaged for the responses generated by GPT-4 **(Table S3)**.

## Acknowledgments

This work was supported by core services from the Cancer Center Support Grant of the Rutgers Cancer Institute of New Jersey (P30CA072720), by the National Institutes of Health (R01CA226537 to R.P. and W.A.), and by the Levy-Longenbaugh Donor-Advised Fund (to R.P. and W.A.). RCSB Protein Data Bank is jointly funded by the National Science Foundation (DBI-1832184, PI: S.K. Burley), the US Department of Energy (DE-SC0019749, PI: S.K. Burley), and the National Cancer Institute, the National Institute of Allergy and Infectious Diseases, and the National Institute of General Medical Sciences of the National Institutes of Health (R01GM133198, PI: S.K. Burley). Molecular graphics and analyses performed with UCSF ChimeraX, developed by the Resource for Biocomputing, Visualization, and Informatics at the University of California, San Francisco, with support from National Institutes of Health R01-GM129325 and the Office of Cyber Infrastructure and Computational Biology, National Institute of Allergy and Infectious Diseases.

## Author Contributions

A.M.I. conceptualization; A.M.I., C.M., and S.K.B. methodology; A.M.I. and C.M. investigation; A.M.I. data curation; A.M.I., C.M., S.K.B., M.B.M., R.P., and W.A. formal analysis; A.M.I. and C.M. writing - original draft; S.K.B., M.B.M., R.P., and W.A. writing - review & editing; S.K.B., R.P., and W.A. funding acquisition; R.P. and W.A. overall project supervision.

