## Supplementary Material for "Generative artificial intelligence performs rudimentary structural biology modeling"

Alexander M. Ille et al.

**Table S1. Exemplary dialog with GPT-4 for amino acid modeling**

| | |
|---|---|
| **User Prompt** | What are the typical distances and angles between the atoms of one arginine residue in a protein? Based on these values, generate a structure in PDB file format for one arginine residue. Ensure coordinate values have three decimal places and omit hydrogen atoms. |
| **GPT-4 Response** | To generate a PDB file format for an arginine residue, we'll need to consider the typical bond lengths and angles found in proteins for this amino acid. The structure of arginine consists of a five-membered aliphatic chain ending in a guanidinium group. The key bonds and angles we need to consider are those in the backbone (common to all amino acids) and those specific to the arginine side chain.<br>Here are typical bond lengths and angles in protein structures:<br>1. Peptide Backbone (common to all amino acids):<br>• N-Cα bond length: ~1.46 Å<br>• Cα-C bond length: ~1.52 Å<br>• N-Cα-C bond angle: ~110°<br>2. Arginine Side Chain:<br>• Cα-Cβ bond length: ~1.53 Å<br>• Cβ-Cγ bond length: ~1.52 Å<br>• Cγ-Cδ bond length: ~1.51 Å<br>• Cδ-Nε bond length: ~1.33 Å<br>• Nε-Cζ bond length: ~1.33 Å<br>• Cζ-Nη1/Nη2 bond length: ~1.33 Å<br>• Bond angles around the aliphatic chain are typically ~109.5° (tetrahedral).<br>• Guanidinium group (Cζ-Nη1/Nη2) angles are typically ~120°.<br>To simplify, we'll not include the hydrogen atoms and consider only the backbone and side chain atoms up to the guanidinium group. We'll assume idealized positions based on typical bond lengths and angles. The PDB format includes columns that are strictly formatted, with atom names, residue names, chain identifiers, residue sequence numbers, and x, y, z coordinates.<br>Here's a simplified PDB file representation of an arginine residue: |
| | CSS |
|  | `ATOM 1 N ARG A 1 0.000 0.000 0.000`<br>`ATOM 2 CA ARG A 1 1.460 0.000 0.000`<br>`ATOM 3 C ARG A 1 2.160 1.370 0.000`<br>`ATOM 4 O ARG A 1 2.160 2.170 -1.080`<br>`ATOM 5 CB ARG A 1 2.090 -1.180 -0.670`<br>`ATOM 6 CG ARG A 1 2.790 -2.440 0.110`<br>`ATOM 7 CD ARG A 1 3.420 -3.620 -0.560`<br>`ATOM 8 NE ARG A 1 4.120 -4.880 0.220`<br>`ATOM 9 CZ ARG A 1 4.750 -5.170 1.380`<br>`ATOM 10 NH1 ARG A 1 4.750 -4.450 2.490`<br>`ATOM 11 NH2 ARG A 1 5.350 -6.340 1.550` |
| | This representation is very simplified and assumes idealized bond lengths and angles. The coordinates (x, y, z) are in angstroms (Å). The atom names (N, CA, C, O, CB, CG, CD, NE, CZ, NH1, NH2) follow the standard nomenclature. The residue name is "ARG" for arginine. The chain identifier is "A", and the residue sequence number is "1".<br>In a real protein structure, the exact coordinates would differ due to the influence of the protein's overall 3D structure, interactions between residues, and flexibility in bond angles and lengths. |

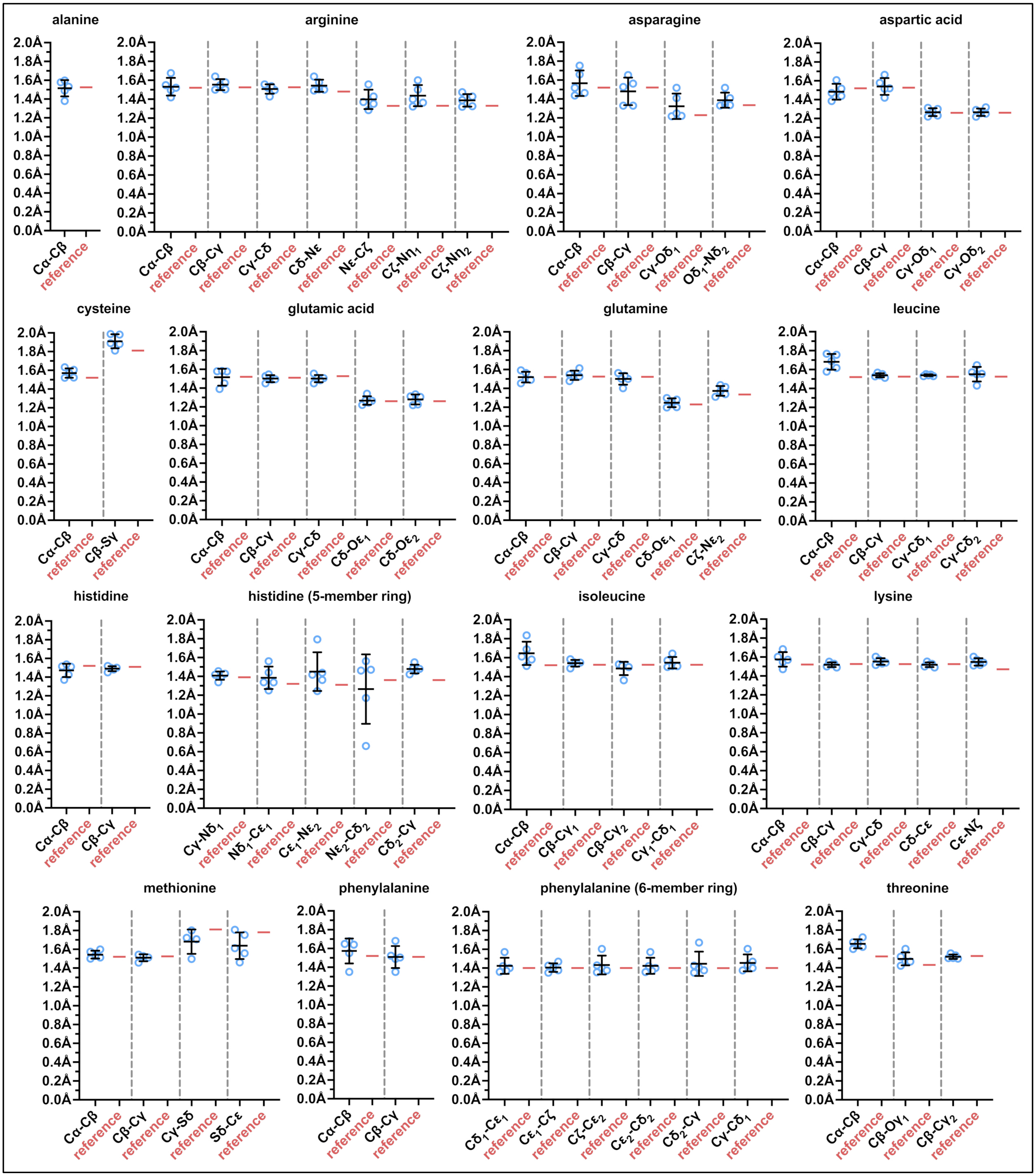


**Fig. S1 Sidechain bond lengths of amino acid structures modeled by GPT-4.** Data shown as means ± SD, n=5 per amino acid (n=4 for methionine bond lengths involving the Cγ atom since one of the methionine modeling iterations resulted in a structure which lacked this atom).

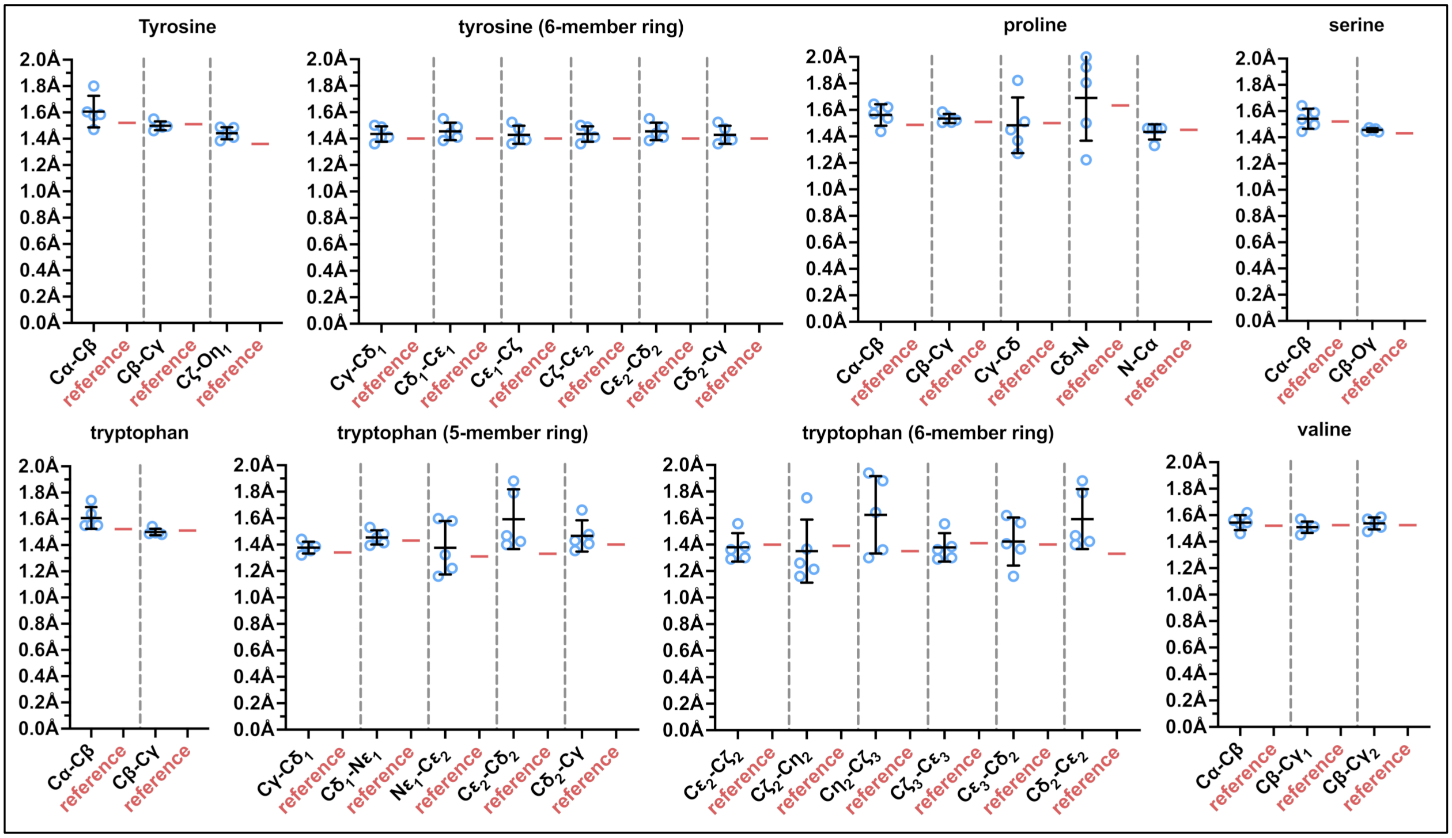


**Fig. S2 Sidechain bond lengths of amino acid structures modeled by GPT-4 (continued)**. Data shown as means ± SD, n=5 per amino acid.

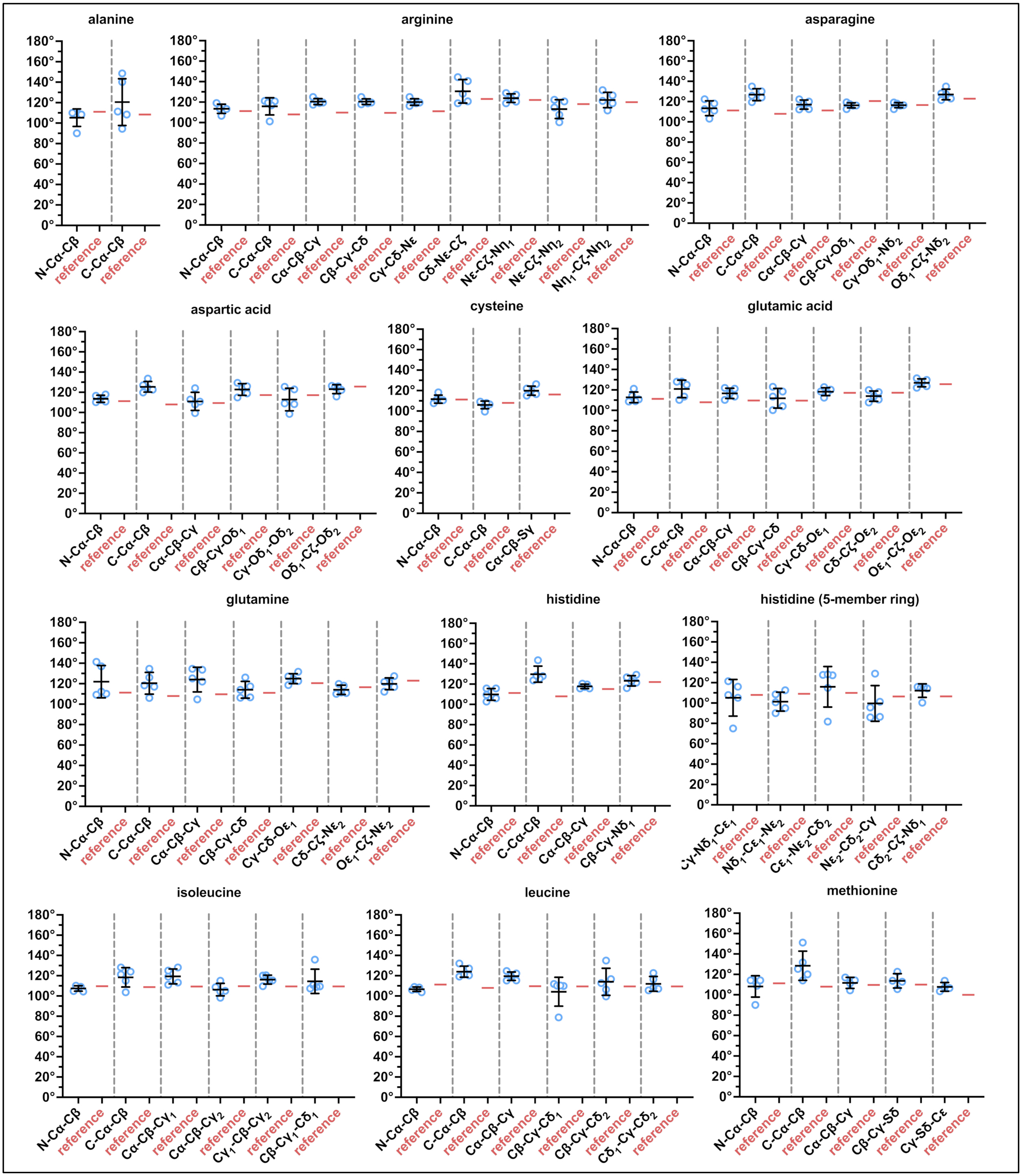


**Fig. S3 Sidechain bond angles of amino acid structures modeled by GPT-4.** Data shown as means ± SD, n=5 per amino acid (n=4 for methionine bond angles involving the Cγ atom since one of the methionine modeling iterations resulted in a structure which lacked this atom).

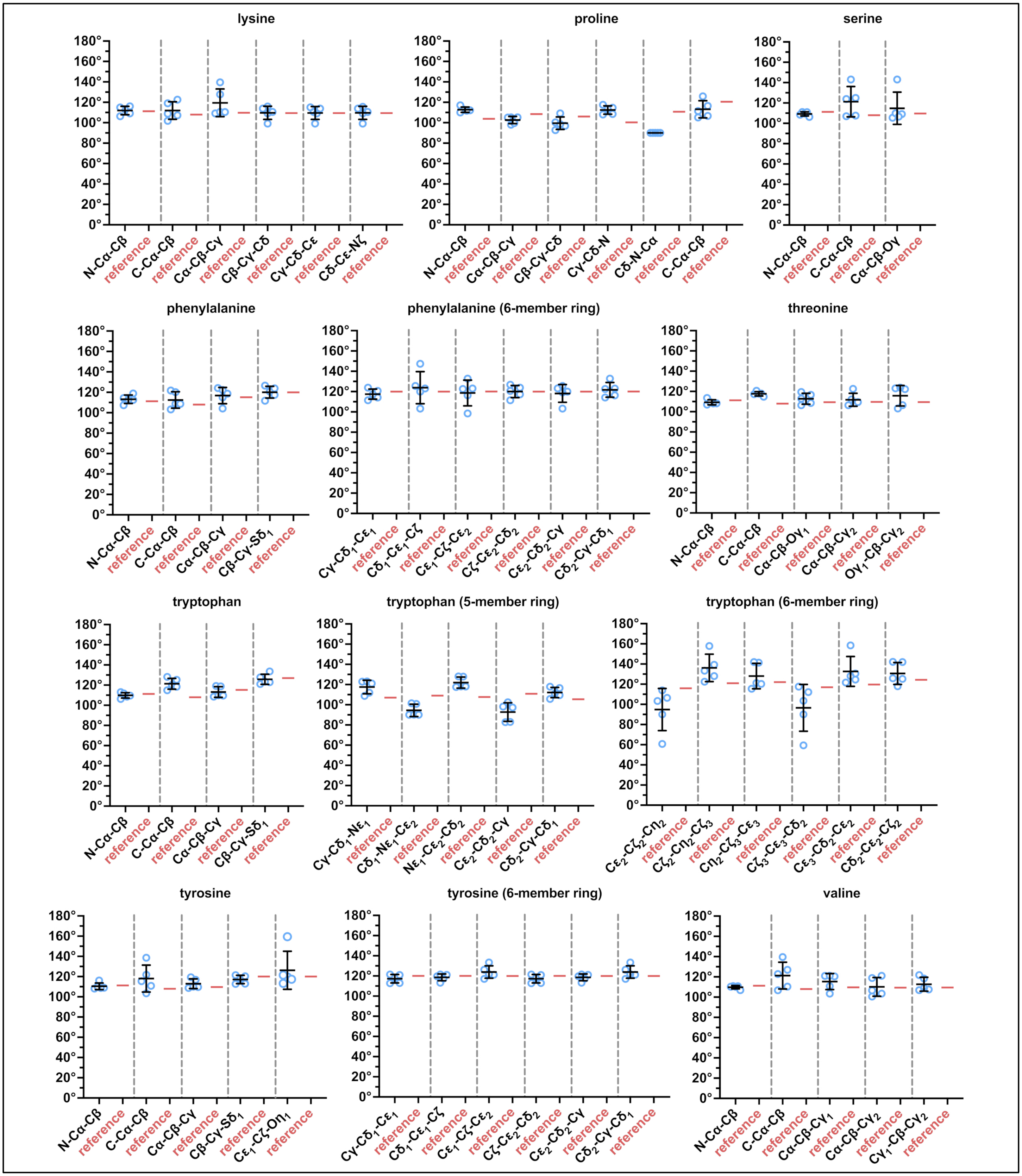


**Fig. S4 Sidechain bond angles of amino acid structures modeled by GPT-4 (continued)**. Data shown as means ± SD, n=5 per amino acid.

**Table S2. Exemplary dialog with GPT-4 running the Wolfram plugin for α-helix modeling.**

| | |
|---|---|
| **User Prompt** | What are the typical geometric attributes (including distances and angles) between the backbone atoms of a typical alpha-helical polypeptide chain? Based on this information, generate a structure in PDB file format for an alpha-helical polypeptide chain 10 residues in length including only alpha carbon atoms. Ensure coordinate values have 3 decimal places and use the Wolfram plug-in for coordinate calculations but not for PDB file formatting. |
| **GPT-4 Response** | To create a PDB file for an alpha-helical polypeptide chain with 10 residues, we first need to understand the typical geometric attributes of an alpha helix. The key attributes include:<br>1. Rise per Residue: This is the linear distance along the helical axis between consecutive amino acids in the helix. For an alpha helix, this is typically about 1.5 Å (angstroms).<br>2. Rotation per Residue: This is the angle of rotation around the helical axis for each amino acid. In an alpha helix, this is usually about 100 degrees.<br>3. Number of Residues per Turn: An alpha helix typically has 3.6 residues per turn.<br>4. Diameter of the Helix: The diameter of an alpha helix is approximately 5-6 Å.<br>Using these parameters, we can calculate the positions of the alpha carbon (Cα) atoms in a 10-residue alpha helix. The Cα atoms are the central points in the backbone of each amino acid residue.<br>The coordinates of each Cα atom can be calculated using helical geometry. The x, y, and z coordinates for the nth Cα atom can be given by:<br>• $x_n = r\ \cos(\theta_n)$<br>• $y_n = r \sin(\theta_n)$<br>• $z_n = n \times$ rise per residue<br>where $r$ is the radius of the helix, $\theta_n$ is the rotation angle for the nth residue, and the rise per residue is the linear distance along the helical axis between consecutive amino acids.<br>Let's perform these calculations using the Wolfram plugin. We'll assume a helix radius of 2.3 Å, which is typical for an alpha helix. |
| | REQUEST TO WOLFRAM |
| | `{ "input": "helixRadius = 2.3; \nresidues = 10; \nrotationPerResidue = 100 Degree; \nrisePerResidue = 1.5; \ncoordinates = Table[{helixRadius*Cos[n*rotationPerResidue], helixRadius*Sin[n*rotationPerResidue], n*risePerResidue}, {n, 0, residues - 1}];\nN[coordinates, 3]" }` |
| | RESPONSE FROM WOLFRAM |
| | `{ "output": "{{2.3, 0., 0.}, {-0.3993908086339397, 2.265057831928078, 1.5}, {-2.1612930278075893, -0.786646329649038, 3.}, {1.15, -1.9918584287042087, 4.5}, {1.7619022191736493, 1.47841150227904, 6.}, {-1.7619022191736493, 1.47841150227904, 7.5}, {-1.15, -1.9918584287042087, 9.}, {2.1612930278075893, -0.786646329649038, 10.5}, {0.3993908086339397, 2.265057831928078, 12.}, {-2.3, 0., 13.5}}" }` |
| | The calculated coordinates for the alpha carbon atoms in a 10-residue alpha helix are as follows (in Ångstroms):<br>1. (2.3, 0.0, 0.0)<br>2. (-0.399, 2.265, 1.5)<br>3. (-2.161, -0.787, 3.0)<br>4. (1.15, -1.992, 4.5)<br>5. (1.762,1.478, 6.0)<br>6. (-1.762, 1.478, 7.5)<br>7. (-1.15, -1.992, 9.0)<br>8. (2.161, -0.787, 10.5)<br>9. (0.399, 2.265, 12.0)<br>10. (-2.3, 0.0, 13.5) |

Now, let's format these coordinates into a PDB file. The PDB file format has a specific structure. Each atom in the structure is represented by an "ATOM" record line, which includes the atom serial number, atom name, residue name, chain identifier, residue sequence number, and the x, y, and z coordinates.
Here is the PDB file content for our 10-residue alpha helix, including only the alpha carbon (Cα) atoms:

CSS

```
ATOM      1  CA  ALA A   1       2.300   0.000   0.000
ATOM      2  CA  ALA A   2      -0.399   2.265   1.500
ATOM      3  CA  ALA A   3      -2.161  -0.787   3.000
ATOM      4  CA  ALA A   4       1.150  -1.992   4.500
ATOM      5  CA  ALA A   5       1.762   1.478   6.000
ATOM      6  CA  ALA A   6      -1.762   1.478   7.500
ATOM      7  CA  ALA A   7      -1.150  -1.992   9.000
ATOM      8  CA  ALA A   8       2.161  -0.787  10.500
ATOM      9  CA  ALA A   9       0.399   2.265  12.000
ATOM     10  CA  ALA A  10      -2.300   0.000  13.500
END
```

In this file, each line represents an alpha carbon atom. "ALA" stands for alanine, a common amino acid, used here to represent each residue. The chain identifier is "A", and the residue sequence numbers run from 1 to 10. The coordinates are aligned in columns as per PDB format standards.

**Table S3. Dialog with GPT-4 for structural drug interaction analysis**

**User Prompt**

Based on structural information, what is the ligand that is present in the attached PDB file? [attached file: 7vh8.pdb]

**GPT-4 Response**

The PDB file you've provided is for the crystal structure of the SARS-CoV-2 main protease in complex with the protease inhibitor PF-07321332. This suggests that PF-07321332 is the ligand present in the structure.
To confirm and provide more details, I will now search for the specific ligand information within the file.

ANALYSIS

PYTHON

```
# First, let's read the contents of the PDB file to understand its structure and identify any ligands present.
file_path = '/mnt/data/7vh8.pdb'

# Open and read the file
with open(file_path, 'r') as file:
    pdb_content = file.readlines()

# Display the first few lines of the file for inspection
pdb_content[:10]
```

RESULT

```
['HEADER    VIRAL PROTEIN                           21-SEP-21   7VH8              \n',
 'TITLE     CRYSTAL STRUCTURE OF SARS-COV-2 MAIN PROTEASE IN COMPLEX WITH PROTEASE\n',
 'TITLE    2 INHIBITOR PF-07321332                                                \n',
 'COMPND    MOL_ID: 1;                                                            \n',
 'COMPND   2 MOLECULE: 3C-LIKE PROTEINASE;                                        \n',
 'COMPND   3 CHAIN: A;                                                            \n',
 'COMPND   4 SYNONYM: MAIN PROTEASE;                                              \n',
 'COMPND   5 EC: 3.4.22.69;                                                       \n',
 'COMPND   6 ENGINEERED: YES                                                      \n',
 'SOURCE    MOL_ID: 1;                                                            \n']
```

In the PDB file, there are entries for ligands indicated by both "HETATM" and "HET" records. Two distinct ligands are identified:

1. A ligand with the identifier "4WI" in chain A, assigned the residue number 401.
2. Another ligand denoted as "DMS" in chain A, with the residue number 402.

The "HETATM" records provide detailed atomic positions for these ligands, indicating their presence within the structure. The ligand "4WI" seems to be the primary ligand of interest, given the context provided by the file's header mentioning the protease inhibitor PF-07321332. "DMS" is likely a solvent or a smaller molecule used in the crystallization process, as Dimethyl Sulfoxide (DMS) is commonly used in such contexts.

ANALYSIS

PYTHON

```
# Searching for the ligand information in the PDB file
# Ligands are usually mentioned in the HETATM records of PDB files

ligands = [line for line in pdb_content if line.startswith('HETATM') or
line.startswith('HET ')]

# Display the first few ligand lines for inspection
ligands[:10]
```

RESULT

```
['HET    4WI  A 401      35                                                     \n',
 'HET    DMS  A 402       4                                                     \n',
 'HETATM 2384  C8  4WI A 401     -16.197  21.132 -25.305  0.80 19.70           C  \n',
 'HETATM 2385  C3  4WI A 401     -17.211  19.842 -29.664  0.80 22.25           C  \n',
 'HETATM 2386  C1  4WI A 401     -19.139  17.723 -27.324  0.80 19.65           C  \n',
 'HETATM 2387  C13 4WI A 401     -19.138  16.564 -24.568  0.80 20.96           C  \n',
 'HETATM 2388  C21 4WI A 401     -19.150  14.286 -22.016  0.80 27.34           C  \n',
 'HETATM 2389  C6  4WI A 401     -18.577  21.464 -25.703  0.80 20.54           C  \n',
 'HETATM 2390  C7  4WI A 401     -18.438  22.041 -24.554  0.80 23.78           C  \n',
 'HETATM 2391  C10 4WI A 401     -20.800  15.398 -25.855  0.80 21.39           C  \n']
```

**User Prompt**

Analyze the structure to detect up to 5 amino acid residues in the protein chain which have important bond interactions with the "4WI" ligand without importing any external libraries. List the residues and the bond distances. Based on this information, predict potential mutations in the protein chain which would interfere with binding of the "4WI" ligand.

**GPT-4 Response**

To analyze the structure and detect amino acid residues in the protein chain that have important bond interactions with the "4WI" ligand, we will follow these steps:

1. Identify the Atoms of the "4WI" Ligand: We'll locate all atoms that belong to the "4WI" ligand.
2. Find Neighboring Protein Atoms: For each atom in the "4WI" ligand, we'll search for nearby protein atoms (from amino acid residues) within a typical interaction distance (e.g., hydrogen bonds are typically less than 3.5 Å, and van der Waals interactions are typically considered within a distance of about 4 Å).
3. Identify Unique Amino Acid Residues: We'll identify unique amino acid residues based on the atoms found to be in proximity to the ligand atoms.
4. Calculate Bond Distances: For the identified interactions, we will calculate the distances between the ligand atoms and the nearest protein atoms to confirm the interaction.
5. Predict Potential Mutations: Based on the identified residues and their interactions with the "4WI" ligand, we can predict potential mutations that would likely interfere with the ligand binding by altering the side chain properties (e.g., charge, size, hydrophobicity) that are critical for the interaction.

Let's start by identifying the atoms of the "4WI" ligand and their neighboring protein atoms.

We identified the following amino acid residues in the protein chain that have important bond interactions with the "4WI" ligand, based on their proximity (within 4 Å):

1. CYS A145 - Distance: 1.81 Å
2. HIS A163 - Distance: 2.68 Å
3. GLU A166 - Distance: 2.77 Å
4. HIS A164 - Distance: 3.02 Å
5. THR A190 - Distance: 3.07 Å

These residues are likely crucial for the binding of the "4WI" ligand to the protein. To predict potential mutations that would interfere with the binding of the "4WI" ligand, we would consider changes to these residues that alter the properties necessary for interaction. Here are some potential mutations:

1. CYS A145: Mutation to a residue with a bulkier side chain or different charge (e.g., CYS to ARG) might disrupt the close interaction.
2. HIS A163 and HIS A164: Histidines often play a role in catalysis or binding through their imidazole ring. Mutating to a residue like LEU or ALA would remove this potential interaction.
3. GLU A166: Glutamate's negative charge is important for interactions. Mutation to a neutral or positively charged residue like GLN or LYS could disrupt binding.
4. THR A190: The hydroxyl group of threonine might be involved in hydrogen bonding. Mutation to a residue like VAL would remove this potential interaction.

These mutations could potentially interfere with the binding by altering the steric and electronic complementarity between the protein and the ligand.

ANALYSIS

PYTHON

```
# Extracting coordinates and other details of atoms in the "4WI" ligand
ligand_atoms = [line for line in hetatm_lines if '4WI' in line]

# Parsing out atom details: atom name, x, y, z coordinates
ligand_atom_details = []
for atom in ligand_atoms:
    atom_name = atom[12:16].strip()
    x = float(atom[30:38].strip())
    y = float(atom[38:46].strip())
    z = float(atom[46:54].strip())
    ligand_atom_details.append((atom_name, x, y, z))

ligand_atom_details[:5]  # Displaying the first five atoms for brevity
```

RESULT

```
[('C8', -16.197, 21.132, -25.305),
 ('C3', -17.211, 19.842, -29.664),
 ('C1', -19.139, 17.723, -27.324),
 ('C13', -19.138, 16.564, -24.568),
 ('C21', -19.15, 14.286, -22.016)]
```

**Table S4. Experimentally determined amino acid sidechain bond distance and angle reference values.**

| amino acid | sidechain distances (Å) | sidechain angles (°) |
|---|---|---|
| alanine | Cα-Cβ, 1.525 | N-Cα-Cβ, 111.095; C-Cα-Cβ, 108.327 |
| arginine | Cα-Cβ, 1.520; Cβ-Cγ, 1.525; Cγ-Cδ, 1.526; Cδ-Nε, 1.480; Nε-Cζ, 1.329; Cζ-Nη$_1$, 1.330; Cζ-Nη$_2$, 1.330 | N-Cα-Cβ, 111.259; C-Cα-Cβ, 107.936; Cα-Cβ-Cγ, 109.705; Cβ-Cγ-Cδ, 109.484; Cγ-Cδ-Nε, 111.011; Cδ-Nε-Cζ, 123.019; Nε-Cζ-Nη$_1$, ;122.022 Nε-Cζ-Nη$_2$, 118.005; Nη$_1$-Cζ-Nη$_2$, 119.973 |
| asparagine | Cα-Cβ, 1.521; Cβ-Cγ, 1.522; Cγ-Oδ$_1$, 1.229; Oδ$_1$-Nδ$_2$, 1.335 | N-Cα-Cβ, 111.288; C-Cα-Cβ, 107.933; Cα-Cβ-Cγ, 111.289; Cβ-Cγ-Oδ$_1$, 120.513; Cγ-Oδ$_1$-Nδ2, 116.612; Oδ$_1$-Cζ-Nδ$_2$, 122.874 |
| aspartic acid | Cα-Cβ, 1.521; Cβ-Cγ, 1.527; Cγ-Oδ$_1$, 1.259; Cγ-Oδ$_2$, 1.260 | N-Cα-Cβ, 111.270; C-Cα-Cβ, 107.916; Cα-Cβ-Cγ, 109.310; Cβ-Cγ-Oδ1, 117.206; Cγ-Oδ$_1$-Oδ$_2$, 117.193; Oδ$_1$-Cζ-Oδ$_2$, 125.601 |
| cysteine | Cα-Cβ, 1.521; Cβ-Sγ,1.810 | N-Cα-Cβ, 111.249; C-Cα-Cβ, 107.952; Cα-Cβ-Sγ, 116.185 |
| glutamic acid | Cα-Cβ, 1.521; Cβ-Cγ, 1.510; Cγ-Cδ, 1.527; Cδ-Oε$_1$, 1.260; Cδ-Oε$_2$, 1.260 | N-Cα-Cβ, 111.288; C-Cα-Cβ, 107.914; Cα-Cβ-Cγ, 109.689; Cβ-Cγ-Cδ, 109.511; Cγ-Cδ-Oε$_1$, 117.157; Cδ-Cζ-Oε$_2$, 117.223; Oε$_1$-Cζ-Oε$_2$, 125.620 |
| glutamine | Cα-Cβ, 1.521; Cβ-Cγ, 1.525; Cγ-Cδ, 1.523; Cδ-Oε$_1$, 1.229; Cζ-Nε$_2$, 1.335 | N-Cα-Cβ, 111.249; C-Cα-Cβ, 107.952; Cα-Cβ-Cγ, 109.639; Cβ-Cγ-Cδ, 111.096; Cγ- |

| | | |
|---|---|---|
| | | Cδ-O$\varepsilon_1$, 120.503; Cδ-Cζ-N$\varepsilon_2$, 116.570; O$\varepsilon_1$-Cζ-N$\varepsilon_2$, 122.927 |
| glycine | N/A | N/A |
| histidine | Cα-Cβ, 1.521; Cβ-Cγ, 1.509; Cγ-N$\delta_1$, 1.391; N$\delta_1$-C$\varepsilon_1$, 1.320; C$\varepsilon_1$-N$\varepsilon_2$, 1.310; N$\varepsilon_2$-C$\delta_2$, 1.360; C$\delta_2$-Cγ, 1.360 | N-Cα-Cβ, 111.270; C-Cα-Cβ, 107.916; Cα-Cβ-Cγ, 115.184; Cβ-Cγ-N$\delta_1$, 122.014; Cγ-N$\delta_1$-C$\varepsilon_1$, 107.989; N$\delta_1$-C$\varepsilon_1$-N$\varepsilon_2$, 109.013; C$\varepsilon_1$-N$\varepsilon_2$-C$\delta_2$, 109.979; N$\varepsilon_2$-C$\delta_2$-Cγ, 106.429; C$\delta_2$-Cζ-N$\delta_1$, 106.591 |
| isoleucine | Cα-Cβ, 1.521; Cβ-C$\gamma_1$, 1.525; Cβ-C$\gamma_2$, 1.525; C$\gamma_1$-C$\delta_1$, 1.525 | N-Cα-Cβ, 109.636; C-Cα-Cβ, 108.777; Cα-Cβ-C$\gamma_1$, 109.365; Cα-Cβ-C$\gamma_2$, 109.641; C$\gamma_1$-Cβ-C$\gamma_2$, 109.478; Cβ-C$\gamma_1$-C$\delta_1$, 109.454 |
| leucine | Cα-Cβ, 1.521; Cβ-Cγ, 1.525; Cγ-C$\delta_1$, 1.524; Cγ-C$\delta_2$, 1.525 | N-Cα-Cβ, 111.262; C-Cα-Cβ, 107.916; Cα-Cβ-Cγ, 109.676; Cβ-Cγ-C$\delta_1$, 109.438; Cβ-Cγ-C$\delta_2$, 109.460; C$\delta_1$-Cγ-C$\delta_2$, 109.480 |
| lysine | Cα-Cβ, 1.521; Cβ-Cγ, 1.526; Cγ-Cδ, 1.525; Cδ-Cε, 1.525; Cε-Nζ, 1.470 | N-Cα-Cβ, 111.220; C-Cα-Cβ, 107.974; Cα-Cβ-Cγ, 109.708; Cβ-Cγ-Cδ, 109.461; Cγ-Cδ-Cε, 109.433; Cδ-Cε-Nζ, 109.469 |
| methionine | Cα-Cβ, 1.521; Cβ-Cγ, 1.525; Cγ-Sδ, 1.811; Sδ-Cε, 1.781 | N-Cα-Cβ,111.288 ; C-Cα-Cβ, 107.914; Cα-Cβ-Cγ, 109.656; Cβ-Cγ-Sδ, 109.990; Cγ-Sδ-Cε, 99.981 |
| phenylalanine | Cα-Cβ, 1.521; Cβ-Cγ, 1.511; Cγ-C$\delta_1$, 1.400; C$\delta_1$-C$\varepsilon_1$, 1,400; C$\varepsilon_1$-Cζ, 1.400; Cζ-C$\varepsilon_2$, 1.400; C$\varepsilon_2$-C$\delta_2$, 1.401; C$\delta_2$-Cγ, 1.399 | N-Cα-Cβ, 111.249; C-Cα-Cβ, 107.971; Cα-Cβ-Cγ, 115.183; Cβ-Cγ-S$\delta_1$, 119.971; Cγ-C$\delta_1$-C$\varepsilon_1$, 119.993; C$\delta_1$-C$\varepsilon_1$-Cζ, 119.968; C$\varepsilon_1$-Cζ-C$\varepsilon_2$, 120.033; Cζ-C$\varepsilon_2$-C$\delta_2$, 119.980; C$\varepsilon_2$-C$\delta_2$-Cγ, 119.987; C$\delta_2$-Cγ-C$\delta_1$, 120.040 |
| proline | Cα-Cβ, 1.487; Cβ-Cγ, 1.509; Cγ-Cδ, 1.500; Cδ-N, 1.639; N-Cα, 1.449 | N-Cα-Cβ, 103.890; Cα-Cβ-Cγ, 108.446; Cβ-Cγ-Cδ, 106.027; Cγ-Cδ-N, 100.384; Cδ-N-Cα, 110.748; C-Cα-Cβ, 120.565 |
| serine | Cα-Cβ, 1.520; Cβ-Oγ, 1.430 | N-Cα-Cβ, 111.259; C-Cα-Cβ, 107.936; Cα-Cβ-Oγ, 109.681 |
| threonine | Cα-Cβ, 1.521; Cβ-O$\gamma_1$, 1.430; Cβ-C$\gamma_2$, 1.525 | N-Cα-Cβ, 111.220; C-Cα-Cβ, 107.974; Cα-Cβ-O$\gamma_1$, 109.328; Cα-Cβ-C$\gamma_2$, 109.696; O$\gamma_1$-Cβ-C$\gamma_2$, 109.419 |
| tryptophan | Cα-Cβ, 1.522; Cβ-Cγ, 1.510; Cγ-C$\delta_1$, 1.340; C$\delta_1$-N$\varepsilon_1$, 1.430; N$\varepsilon_1$-C$\varepsilon_2$, 1.311; C$\varepsilon_2$-C$\delta_2$, 1.330; C$\delta_2$-Cγ, 1.400; C$\varepsilon_2$-C$\zeta_2$, 1.399; C$\zeta_2$-C$\eta_2$, 1.390; C$\eta_2$-C$\zeta_3$, 1.350; C$\zeta_3$-C$\varepsilon_3$, 1.411; C$\varepsilon_3$-C$\delta_2$, 1.401; C$\delta_2$-C$\varepsilon_2$, 1.330 | N-Cα-Cβ, 111.276; C-Cα-Cβ, 107.904; Cα-Cβ-Cγ, 115.228; Cβ-Cγ-S$\delta_1$, 126.989; Cγ-C$\delta_1$-N$\varepsilon_1$, 107.014; C$\delta_1$-N$\varepsilon_1$-C$\varepsilon_2$, 109.000; N$\varepsilon_1$-C$\varepsilon_2$-C$\delta_2$, 107.630; C$\varepsilon_2$-C$\delta_2$-Cγ, 110.881; C$\delta_2$-Cγ-C$\delta_1$, 105.476; C$\varepsilon_2$-C$\zeta_2$-C$\eta_2$, 115.995; C$\zeta_2$-C$\eta_2$-C$\zeta_3$, 120.991; C$\eta_2$-C$\zeta_3$-C$\varepsilon_3$, 122.039; C$\zeta_3$-C$\varepsilon_3$-C$\delta_2$, 116.940; C$\varepsilon_3$-C$\delta_2$-C$\varepsilon_2$, 119.677; C$\delta_2$-C$\varepsilon_2$-C$\zeta_2$, 124.356 |
| tyrosine | Cα-Cβ, 1.521; Cβ-Cγ, 1.510; Cζ-O$\eta_1$, 1.360; Cγ-C$\delta_1$, 1.400; C$\delta_1$-C$\varepsilon_1$, 1.400; C$\varepsilon_1$-Cζ, 1.400; Cζ-C$\varepsilon_2$, 1.400; C$\varepsilon_2$-C$\delta_2$, 1.400; C$\delta_2$-Cγ, 1.401 | N-Cα-Cβ, 111.220; C-Cα-Cβ, 107.974; Cα-Cβ-Cγ, 109.697; Cβ-Cγ-S$\delta_1$, 120.052; C$\varepsilon_1$-Cζ-O$\eta_1$, 120.014; Cγ-C$\delta_1$-C$\varepsilon_1$, 120.010; C$\delta_1$-C$\varepsilon_1$-Cζ, 120.032; C$\varepsilon_1$-Cζ-C$\varepsilon_2$, 119.988; Cζ-C$\varepsilon_2$-C$\delta_2$, 120.015; C$\varepsilon_2$-C$\delta_2$-Cγ, 119.997; C$\delta_2$-Cγ-C$\delta_1$, 119.958 |
| valine | Cα-Cβ, 1.521; Cβ-C$\gamma_1$, 1.525; Cβ-C$\gamma_2$, 1.525 | N-Cα-Cβ, 111.270; C-Cα-Cβ, 107.916; Cα-Cβ-C$\gamma_1$, 109.673; Cα-Cβ-C$\gamma_2$, 109.390; C$\gamma_1$-Cβ-C$\gamma_2$, 109.490 |

**Table S5. Sequence used as input for AlphaFold2 modeling.** In order to meet minimum input requirements, the sequence was composed of 10 proline residues followed by 30 alanine residues. From the modeled structure, only alanine residues from positions 21-30 were used as the α-helix structure for analysis.

| **proline stretch** | **alanine stretch** | | |
|---|---|---|---|
| positions 1-10 | positions 11-20 | positions 21-30 | positions 31-40 |
| PPPPPPPPPP | AAAAAAAAAA | AAAAAAAAAA | AAAAAAAAAA |

**Fig. S5 Structures of α-helices modeled with AlphaFold2 (A), ChimeraX (B), and PyMOL (C).**

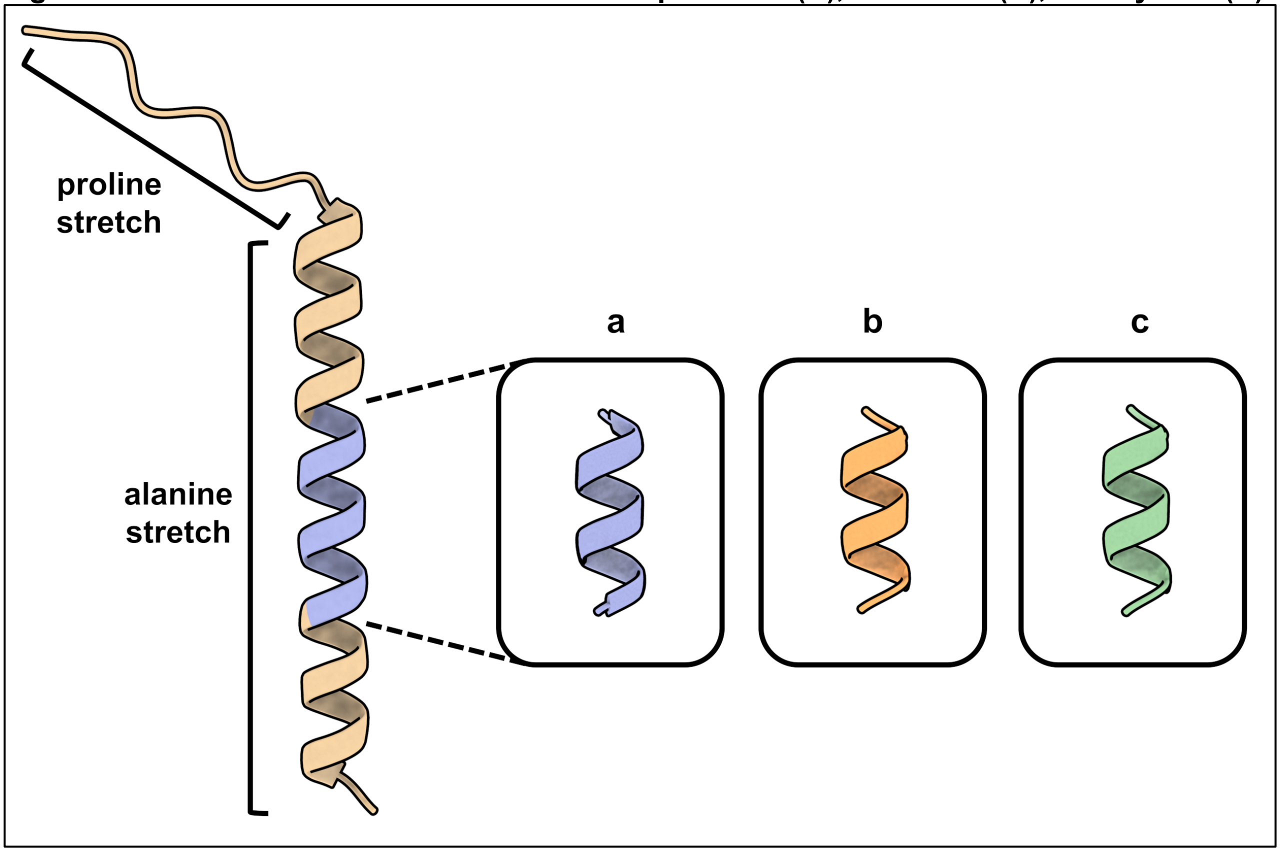
